# Biallelic mutation of *CLRN2* causes non-syndromic hearing loss in humans

**DOI:** 10.1101/2020.07.29.222828

**Authors:** Barbara Vona, Neda Mazaheri, Sheng-Jia Lin, Lucy A Dunbar, Reza Maroofian, Hela Azaiez, Kevin T. Booth, Sandrine Vitry, Aboulfazl Rad, Pratishtha Varshney, Ben Fowler, Christian Beetz, Kumar N. Alagramam, David Murphy, Gholamreza Shariati, Alireza Sedaghat, Henry Houlden, Shruthi VijayKumar, Richard J. H. Smith, Thomas Haaf, Aziz El-Amraoui, Michael R. Bowl, Gaurav K. Varshney, Hamid Galehdari

## Abstract

Deafness, the most frequent sensory deficit in humans, is extremely heterogenous with hundreds of genes probably involved. Clinical and genetic analyses of an extended consanguineous family with pre-lingual, moderate-to-profound autosomal recessive sensorineural hearing loss, allowed us to identify *CLRN2*, encoding a tetraspan protein as a new deafness gene. Homozygosity mapping followed by exome sequencing identified a 15.2 Mb locus on chromosome 4p15.32p15.1 containing a missense pathogenic variant in *CLRN2* (c.494C>A, NM_001079827.2) segregating with the disease. Using *in vitro* RNA splicing analysis, we show that the *CLRN2* c.494C>A mutation leads to two events: 1) the substitution of a highly conserved threonine (uncharged amino acid) to lysine (charged amino acid) at position 165, p.(Thr165Lys), and 2) aberrant splicing, with the retention of intron 2 resulting in a stop codon after 26 additional amino acids, p.(Gly146Lysfs*26). Expression studies and phenotyping of newly produced zebrafish and mouse models deficient for clarin 2 further confirm that clarin 2, expressed in the inner ear hair cells, is essential for normal organization and maintenance of the auditory hair bundles, and for hearing function. Together, our findings identify *CLRN2* as a new deafness gene, which will impact future diagnosis and treatment for deaf patients.

## Introduction

The mammalian inner ear is an exquisite and highly complex organ, made up of the vestibule, the organ responsible for balance, and the cochlea, the sensory organ for hearing. The auditory sensory cells of the inner ear are called the inner and outer hair cells that are responsible for transduction of sound wave-induced mechanical energy into neuronal signals (*1, 2*). The functional mechanoelectrical transduction machinery involves intact formation and maintenance of a highly specialized and organized structure, the hair bundle. The hair bundle contains a few dozen F-actin-filled stereocilia, arranged in a highly interconnected and highly organized staircase-like pattern, which is critical for function (*3*). Knowledge of the mechanisms of formation, maintenance, and function of the transduction complex is limited (*4*). In this regard, identification of novel genes that encode protein products essential for hearing is likely to improve our understanding of the physical, morphological and molecular properties of hair cells and associated mechanistic processes.

Hereditary hearing loss is one of the most common and genetically heterogeneous disorders in humans (*5*). Sensorineural hearing loss has an incidence of one to two per 1000 at birth (*6*). It displays extraordinary phenotypic, genetic and allelic heterogeneity, with up to 1,000 different genes potentially involved (*7*). So far, about 120 genes and more than 6,000 disease causing variants (*8*) have been identified as responsible for non-syndromic hearing loss in humans (see http://hereditaryhearingloss.org/ and http://deafnessvariationdatabase.org/), and many more are yet to be discovered. Genetic factors predominate the etiological spectrum and most of hereditary hearing loss appears to follow an autosomal recessive inheritance pattern (*9*). Approximately 80% of the currently known autosomal recessive genes have been identified by studying extended consanguineous families (*10*). There are many forms of hearing loss that are clinically indistinguishable but caused by distinct genetic entities that are presently unknown. Identification of additional genes essential for auditory function, through the study of families exhibiting hereditary hearing loss, will not only help increase our understanding of the biology of hearing, but will also identify new molecular targets for therapeutic intervention.

Through the study of an extended consanguineous Iranian family, we have identified a *CLRN2* coding lesion as the cause of hearing loss in family members that are homozygous for the allele. We have established that clarin 2 likely plays a critical role in mechanotransducing stereocilia of the hair bundle in zebrafish and mouse. *CLRN2* belongs to the clarin (CLRN) family of proteins that are comprised of three orthologues named clarin 1, 2, and 3 that encode four-transmembrane domain proteins. Pathogenic variants in *CLRN1* (clarin 1) cause either non-syndromic retinitis pigmentosa (RP) (*11*) or Usher syndrome type 3A (USH3A), that is characterized by progressive hearing loss, RP and variable vestibular dysfunction (*12-15*). This study establishes clarin 2 as essential for inner ear function in zebrafish, mice and humans, with a loss-of-function allele leading to autosomal recessive non-syndromic sensorineural hearing loss (ARNSHL).

## Materials and Methods

### Patient clinical and audiometry data

A three generation Iranian family of Lurs ethnicity was ascertained as part of a large ethnically diverse Iranian population rare disease study. After obtaining written informed consent from all participants with approval by the Faculty of Medicine ethics commissions at the University of Würzburg (46/15) and Shahid Chamran University of Ahvaz (#EE/97.24.3 17654), pure-tone audiograms and medical information were collected from participating members. Clinical examination excluded additional syndromic features.

Individuals IV-1, IV-6, and V-1 (Figure 1) underwent complete ear, nose and throat examination, including binocular ear microscopy and external ear inspection. Routine pure-tone audiometry was performed according to current standards that measured hearing thresholds at frequencies 0.25, 0.5, 1, 2, 4, 6 and 8 kHz. Both air-and bone-conduction thresholds were determined. Severity of hearing loss was defined as previously described (*16*). Individuals IV-1 and IV-6 underwent additional tympanometry and speech recognition threshold testing. Audiometry testing for individuals IV-1, IV-6, and V-1 was performed at ages 29 years, 44 years, and 20 years, respectively.

**Fig. 1.**
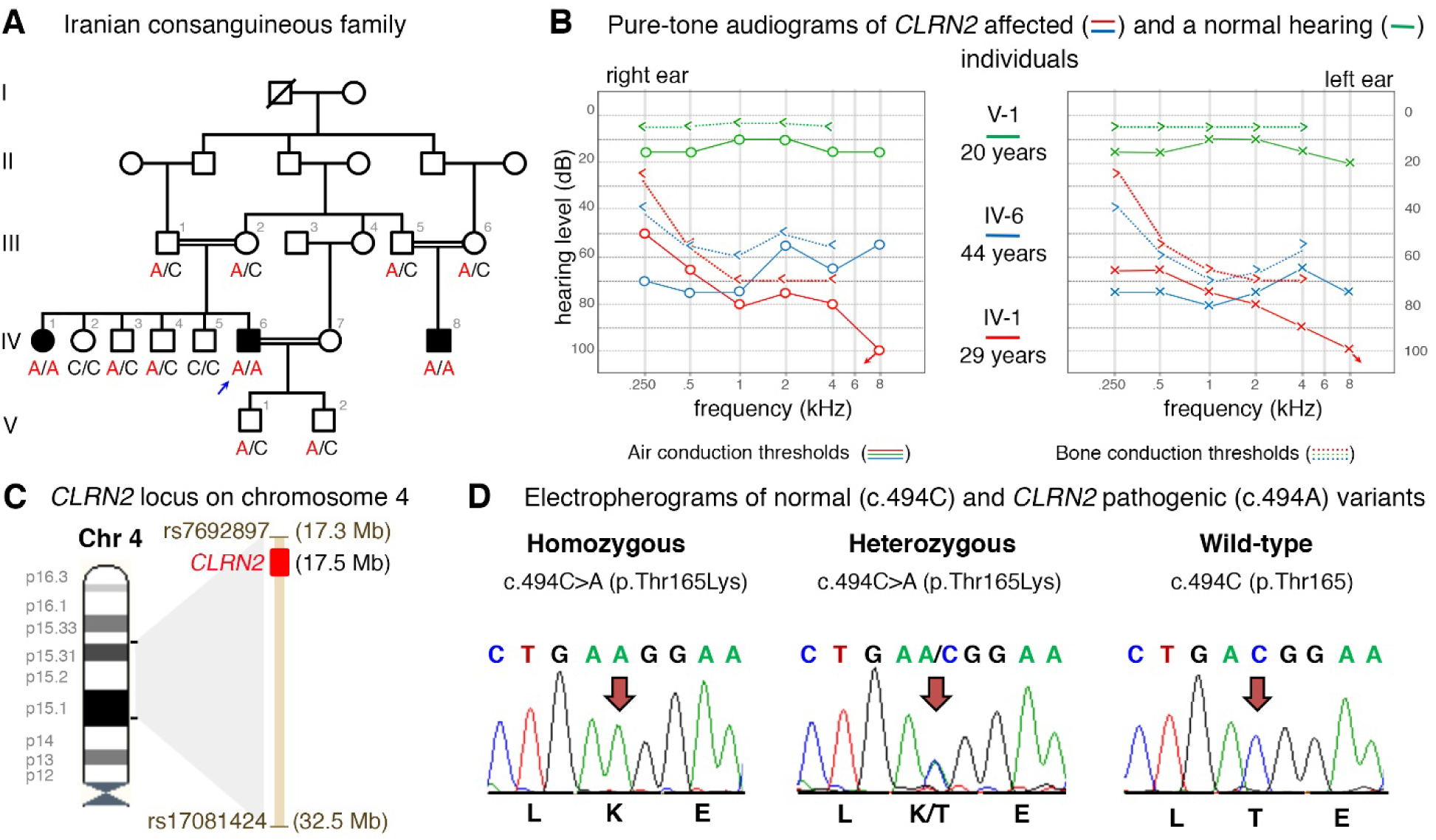
Pedigree, audiological data, genetic data, and locus mapping. **(A)** The consanguineous family of Iranian origin with hearing loss and segregation of the *CLRN2* c.494C>A variant. **(B)** Pure-tone audiograms from affected individuals IV-1 (red) and IV-6 (blue), as well as an unaffected heterozygous individual V-1 (green). Air conduction thresholds in dB HL for the right and left ears are represented by circles and crosses, respectively. Bone-conduction thresholds are represented by < and > for right and left ears, respectively, and confirm a sensorineural hearing loss in the affected individuals. **(C)** Homozygosity mapping reveals a 15.2 Mb locus on chromosome 4 containing *CLRN2*. **(D)** Sequence electropherograms showing the homozygous, heterozygous and WT images of the *CLRN2* c.494C>A; pThr165Lys pathogenic variants.

### Genotyping, homozygosity mapping, copy number variation and exome sequencing data analyses

Due to parental consanguinity and suspected autosomal recessive mode of inheritance, we assumed that the causative variant would be homozygous and identical by descent in affected individuals in the fourth generation of the family. Blood samples from 14 family members were obtained and genomic DNA was isolated from whole blood using standard procedures. DNA from affected (IV-1, IV-6, and IV-8) and unaffected (IV-2, IV-3, IV-4, and IV-5) individuals were genotyped using the Infinium Global Screening Array-24 v1.0 BeadChip (Illumina, SanDiego, CA, USA) according to manufacturer’s protocols. Homozygosity mapping was performed using Homozygosity Mapper to identify common homozygous intervals among the affected individuals (*17*). Runs of homozygosity with a maximum threshold of 0.99 were checked after the exome-wide analysis was completed. Copy number variation calling was performed using GenomeStudio v.2011.1 and cnvPartition 3.2.0 (Illumina).

For exome sequencing, DNA samples from two affected individuals (IV-1 and IV-6) were used. The data from individual IV-6 was analyzed exome-wide and data from individual IV-1 was used for determination of allele sharing. Exome capture using genomic DNA was performed using the SureSelect Target Enrichment v6 (Agilent) kit following manufacturer’s recommendations. The libraries were sequenced with a HiSeq4000 (Illumina). Data analysis was performed using the Burrows-Wheeler Alignment (BWA) tool for read mapping to the human reference genome GRCh37 (hg19), Picard for duplicate removal, GATK for local re-alignment, base recalibration, variant calling, and variant annotation, and SnpEff for variant annotation. Variant filtering was based on: coverage > 10X, Phred quality score ≥ 30, and minor allele frequency (MAF) ≤ 0.005 as reported in 1000 Genomes Project and EVS6500. Variants were filtered based on coding effect (non-synonymous, indels, and splice site variants), and artifact-prone genes (*HLA*s, *MAGE*s, *MUC*s, *NBPF*s, *OR*s, *PRAME*s) were excluded. Visualization was performed using the Integrative Genomics Viewer. Analysis of homozygous and compound heterozygous variants between the two sequenced affected individuals (IV-6 and IV-1) followed. We analyzed missense variants by using a combination of criteria that scored conservation using GERP++ and PhyloP, and deleterious or pathogenic scores in Combined Annotation Dependent Depletion (CADD) (*18*), LRT (*19*), MutationTaster (*20*), PolyPhen-2 (*21*), and SIFT (*22*). Missense variants were excluded when three out of five *in silico* pathogenicity prediction tools yielded a benign score. Manual MAF analysis used gnomAD (*23*), GME (*24*), and Iranome (*25*). Potential effects on splicing were assessed using ESEfinder (*26*) and RESCUE-ESE (*27*).

### Segregation, sequence and *in vitro* splicing analyses of the *CLRN2* c.494C>A pathogenic variant

To confirm segregation of the *CLRN2* c.494C>A; p.(Thr165Lys) (NM_001079827.2) homozygous variant, Sanger sequencing was completed in all 14 family members using the following primers (*CLRN2* Ex3 F: 5’-AAATGCCACCTCTTACAGAGTTGC-3’ and *CLRN2* Ex3 R: 5’-ACCGTGGCCTCTTCGATTTTGGTC-3’) and standard PCR and sequencing parameters.

To document residue conservation, CLRN1 (UniProt: P58418) and CLRN2 (UniProt: A0PK11) were aligned and visualized in Jalview (*28*) with an overview of the pathogenic and likely pathogenic missense and nonsense *CLRN1* variants retrieved from the Deafness Variation Database v 8.2 (*8*).

In addition, secondary protein structure prediction of human CLRN2 (NP_001073296.1) that included the wild-type (WT) and mutated amino acid residues was performed using I-TASSER (*29*).

To assess the splicing effect of the c.494C>A variant, *in vitro* splicing assays, also called mini-genes, were carried out as described (*30, 31*). Wild-type (WT) *CLRN2* exon 3 (266 bp) plus 183 and 51 nucleotides from intron 2 and the 3’UTR were PCR amplified with gene-specific primers containing *Sal*l or *Sac*II restriction enzyme sites, respectively. After PCR amplification, clean up, and restriction enzyme digestion, the PCR fragment was ligated into the pET01 Exontrap vector (MoBiTec) and the sequence was confirmed. Variants were then introduced into the WT sequence using QuikChange Lightning Site-Directed Mutagenesis (Agilent) according to the manufacturer’s protocols via overlapping primers containing the alteration. The WT and mutant mini-genes were sequence confirmed.

WT or mutant mini-genes were transfected in triplicates into COS-7 and ARPE-19 cells using TransIT-LT1 Transfection Reagent (Mirus). Cells were harvested 36h after transfection and total RNA was extracted using Quick-RNA MiniPrep Plus kit (ZYMO Research). cDNA was transcribed using 750ng of isolated RNA SuperScript™ III Reverse Transcriptase (ThermoFisher Scientific) using a primer specific to the 3’ native exon of the pET01 vector according to manufacturer’s protocol. PCR amplification followed using primers specific to the 5’ and 3’ native exons of the pET01 vector, and products were visualized on a 1.5% agarose gel. As a negative control, rs117875715 (chr4(GRCh37):g.17,528,480G>A), a benign polymorphism, was used to test and validate the designed mini-gene assay.

Concurrently, the mini-gene splice assay experiment was conducted in a double-blind manner as previously described (*32*). Genomic DNAs of an affected homozygous (IV-6) and WT individual (IV-5) were amplified using a forward primer with a *Xho*I restriction site (*CLRN2* Ex3 *Xho*I F: 5’-aattctcgagTTGCAGTGAGCTGAGATGGT-3’) and a reverse primer with a *Bam*HI restriction site (*CLRN2* Ex3 *Bam*HI R: 5’-attggatccGCCTTGCGAAGTTGTTACTG-3’). The 886 bp amplicon included the entire exon 3 sequence plus additional flanking 320 bp (5’) and 306 bp (3’) sequence that was ligated into a multiple cloning site between native exons A and B in the linearized pSPL3b exon-trapping vector. The vector was transformed into DH5α competent cells and plated overnight. All mutant mini-genes were Sanger sequence confirmed.

Homozygous and WT mini-genes were transfected in triplicate into HEK 293T cells cultured in FCS-free medium in 6 well culture plates with a density of 2×10^5^ cells per mL. The mini-genes in the pSPL3b vector were transiently transfected using 6µl of FuGENE 6 Transfection Reagent (Roche) with 2 µg of vector. An empty vector and HEK 293T cells were included as controls. The transfected cells were harvested 24 to 48 hours post-transfection. Total RNA was prepared using the miRNAeasy Mini Kit (Qiagen). Approximately, 1 µg of RNA was reverse transcribed using a High Capacity RNA-to-cDNA Kit (Applied Biosystems) following manufacturer’s protocols. The cDNA was used for PCR amplification using a vector specific SD6 forward (5’-TCTGAGTCACCTGGACAACC-3’) and a terminal *CLRN2* exon 3 reverse cDNA primer (5’-CAAGATATCCTCAGCTGTGACC-3’). The resulting amplified fragments were visualized on a 1.5% agarose gel. cDNA amplicons were Sanger sequenced. cDNA amplicons from the homozygous individual were cloned following standard protocols for the TA cloning (dual promoter with pCRII) kit (Invitrogen).

### CRISPR/Cas9-mediated inactivation of *clrn2* in zebrafish

Zebrafish (*Danio rerio*) were raised and maintained in an AALAC accredited facility at the Oklahoma Medical Research Foundation (OMRF) under standard conditions. Zebrafish embryos/larvae were maintained in embryo medium with 0.00002% methylene blue and raised at 28°C. All animal experiments were performed as per protocol (17-01) and approved by the Institutional Animal Care Committee of OMRF (IACUC). All zebrafish handling, embryo care, and microinjections were performed as previously described (*33*). WT zebrafish strain NHGRI-1 was used for all experiments (*34*). The zebrafish embryonic staging was determined by morphological features according to Kimmel et al (*35*).

To produce zebrafish *clrn2* crispants, the sgRNA target sequences were selected from the UCSC genome browser tracks generated by the Burgess lab. Five independent targets were chosen and sgRNAs were synthesized by *in vitro* transcription as described earlier (*36*). sgRNAs and Cas9 protein complex were used to generate indels. A 6 µL mixture containing 2 µL of 40 µM Spy Cas9 NLS protein (New England Biolabs, MA, USA), 200 ng each of five sgRNAs (in 2 µL) and 2 µL of 1 M potassium chloride was injected into one-cell-stage WT embryos. Injection volumes were calibrated to 1.4 nL per injection. Insertion/deletion (indel) variants were detected by amplifying the target region by PCR and Sanger sequencing as described earlier (*36*). The sequencing data were analyzed by Inference of CRISPR Edits (ICE) v2 CRISPR analysis tool. The sgRNA target sequences and PCR primer sequences are listed in Table S1.

### Zebrafish RNA extraction and real-time quantitative PCR (RT-qPCR)

Total RNA at different developmental stages, adult tissues, and CRISPR/Cas9 injected larvae were extracted using the TRIzol Reagent (Thermo Fisher Scientific, CA, USA) and purified by RNA clean and concentrator-5 kit (Zymo Research, CA, USA) according to the manufacturer’s instructions. RNA concentration was measured by DeNovix DS-11 spectrophotometer (DeNovix Inc. USA). The cDNA was synthesized by iScript RT Supermix (Bio-Rad, USA), and was used as a template for performing the RT-qPCR with SYBR Green Supermix (Thermo Fisher Scientific, CA, USA) and the Light Cycler® 96 System (Roche, CA, USA). All RT-qPCR reactions were carried out using three biological and technical replicates. The housekeeping gene *18S* was used as a reference gene.

All RT-qPCR primer pairs were designed across exon-exon junctions using NCBI Primer-BLAST program. The sequences are listed in Table S1. The PCR cycling conditions were used as per the manufacturer instructions, and the amplification specificity was assessed by dissociation curve analysis at the end of the PCR cycles. The cycle threshold values (Ct) data were imported into Microsoft Excel for the relative gene expression analysis. Quantification was based on 2^(-ΔΔCT) method (*37*), and using 18 hours post fertilization (hpf) for *clarin 2* temporal expression, muscle for *clarin 2* in different tissue expression and the corresponding age-matched control for *clarin 2* CRISPR injected F_0_ larvae as normalization control.

### Distribution of *clrn2* and phalloidin staining in zebrafish

To determine *clrn2* expression, we used in situ hybridization on larvae and inner ear-containing cryosections. The full-length coding sequence of zebrafish *clarin 2* (NM_001114690.1) was PCR amplified from WT zebrafish cDNA using primer pairs with restriction enzymes *Bam*HI and *Xho*I restriction sites cloned into pCS2+ vector (a kind gift from Dr. Dave Turner, University of Michigan). After restriction digestion, the resulting clones were sequenced and used as templates for riboprobe synthesis. The digoxigenin-UTP-labeled riboprobes were synthesized according to the manufacturer’s instructions (Millipore Sigma, MO, USA). Briefly, the *clarin 2* and the *pvalb9* plasmids (Horizon Discovery) were linearized by *Bam*HI and *Not*I restriction enzymes, respectively. The linearized plasmid was purified and used as template for in vitro transcription using T7 RNA polymerase to synthesize anti-sense probes. The sense probe was made using *Xba*I linearized *clarin 2* plasmid and SP6 RNA polymerase.

The whole-mount in situ hybridization (WISH) on 3 and 5 dpf zebrafish embryos/larvae was performed following the procedures as described by Thisse et al. with minor modifications (*38*). Age-matched zebrafish embryos were randomly collected by breeding WT zebrafish pairs. The embryos were treated with 0.003% phenylthiourea (PTU) (Millipore Sigma, MO, USA) in embryo medium at 1-day post-fertilization (dpf) until the desired stages reached to reduce the pigment formation that will facilitate color visualization during *in situ* hybridization. Embryos/larvae were then fixed with 4% (V/V) paraformaldehyde in phosphate-buffered saline (PBS) at 3 and 5 dpf. An additional bleaching step was carried out after fixation by incubating the embryos at room temperature in a 3% hydrogen peroxide and 0.5% potassium hydroxide solution. The permeabilization of 3 dpf embryos and 5 dpf larvae were using 2 µg/mL proteinase K for 12 and 18 minutes respectively. Color development was conducted using the BM-Purple alkaline phosphatase substrate (Millipore Sigma, MO, USA).

For preparation of cryo-section samples after WISH, the 5 dpf larvae were soaked in 25%, 30% (V/V) sucrose/PBS and optimum cutting temperature (OCT) each for at least two days, and embedded in OCT then Cryotome sectioned at a 10-micrometer thickness.

For phalloidin staining of the zebrafish inner ear, 5 dpf larvae were euthanized with tricaine and fixed in 4% (V/V) paraformaldehyde (PFA) at 5 dpf, fixed embryos were washed by PBSTx (1% PBS, 0.2% triton X-100) and incubated in 2% triton X-100 in PBS at room temperature for overnight with agitation until the otoliths were completely dissolved. The larvae were sequentially washed in PBSTx and incubated with Alexa Fluor 647 Phalloidin (1:100) (Thermo Fisher Scientific, CA, USA) in PBSTw (1% PBS, 0.1% Tween-20) at room temperature for 4 hours. The samples were washed in PBSTx after staining and mounted laterally in 75% glycerol on slides. Images were acquired with a Zeiss LSM-710 Confocal microscope.

### Production and phenotyping of clarin 2 deficient mutant in mice

The *Clrn2*^*del629*^ mutant line was generated on a C57BL/6N background by the Molecular and Cellular Biology group at the Mary Lyon Centre (MLC), MRC Harwell Institute, using CRISPR/Cas9 genome editing (*39*). The mice were housed and maintained under specific pathogen-free conditions in individually ventilated cages, with environmental conditions as outlined in the Home Office Code of Practice. Animals were housed with littermates until weaned, and then housed with mice of the same sex and of similar ages, which was often their littermates. Both male and female animals were used for all experiments. Animal procedures at the MRC Harwell Institute were licenced by the Home Office under the Animals (Scientific Procedures) Act 1986, UK and additionally approved by the Institutional Animal Welfare and Ethical Review Body (AWERB). The *Clrn1−/−* mice (*Clrn1*^tm1.2Ugpa^, MGI: 6099052) used for comparative scanning electron microscopy analyses were previously described (*40*).

Auditory Brainstem Response (ABR) tests were performed using a click stimulus and frequency-specific tone-burst stimuli (at 8, 16 and 32 kHz) to screen mice for auditory phenotypes and investigate auditory function (*41*). Distortion Product Oto-Acoustic Emission (DPOAE) tests were performed using frequency-specific tone-burst stimuli from 8 to 32 kHz with the TDT RZ6 System 3 hardware and BioSig RZ software (Tucker Davis Technology, Alachua, FL, USA). Mice were anaesthetized by intraperitoneal injection of ketamine (100 mg/ml at 10% v/v) and xylazine (20 mg/ml at 5% v/v) administered at the rate of 0.1 ml/10 g body mass. Once fully anaesthetized, mice were placed on a heated mat inside a sound-attenuated chamber (ETS Lindgren) and recording electrodes (Grass Telefactor F-E2-12) were placed subdermally over the vertex (active), right mastoid (reference) and left mastoid (ground). ABRs were collected, amplified and averaged using TDT System 3 hardware and BioSig software (Tucker Davies Technology, Alachua, FL, USA). The click stimulus consisted of a 0.1 ms broadband click presented at a rate of 21.1/s. Tone-burst stimuli were of 7 ms duration including rise/fall gating using a 1 ms Cos2 filter, presented at a rate of 42.5/s. All stimuli were presented free-field to the right ear of the mouse, starting at 90 dB SPL and decreasing in 5 dB increments until a threshold was determined visually by the absence of replicable response peaks. For graphical representation, mice not showing an ABR response at the maximum level tested (90 dB SPL) were recorded as having a threshold of 95 dB SPL. These mice/thresholds were included when calculating genotype average thresholds. All ABRs were performed blind to genotype, to ensure thresholds were obtained in an unbiased manner. Mice were recovered using 0.1 ml of anaesthetic reversal agent atipamezole (Antisedan™, 5 mg/ml at 1% v/v).

DPOAE tests utilized an ER10B+ low noise probe microphone (Etymotic Research) to measure the DPOAE near the tympanic membrane. Tone stimuli were presented via separate MF1 (Tucker Davis Technology) speakers, with f1 and f2 at a ratio of f2/f1 = 1.2 (L1=65 dB SPL, L2=55 dB SPL). Mice were anaesthetized via intraperitoneal injection of ketamine (100 mg/ml at 10% v/v), xylazine (20 mg/ml at 5% v/v) and acepromazine (2 mg/ml at 8% v/v) administered at a rate of 0.1 ml/10 g body mass. Once surgical anaesthesia was confirmed by the absence of a pedal reflex, a section of the pinna was removed to enable unobstructed access to the external auditory meatus. Mice were then placed on a heated mat inside a sound-attenuated chamber (ETS-Lindgren) and a pipette tip containing the DPOAE probe assembly was inserted into the ear canal. In-ear calibration was performed before each test. The f1 and f2 tones were presented continuously and a fast-Fourier transform was performed on the averaged response of 356 epochs (each ∼21 ms). The level of the 2f1-f2 DPOAE response was recorded and the noise floor calculated by averaging the four frequency bins either side of the 2f1-f2 frequency.

Mice were euthanized by cervical dislocation and inner ears were removed and fixed in 2.5% glutaraldehyde (TAAB Laboratories Equipment Ltd.) in 0.1 M phosphate buffer (Sigma-Aldrich) overnight at 4°C. Following decalcification in 4.3% EDTA, cochleae were sub-dissected to expose the sensory epithelium then ‘OTO processed’ with alternating incubations in 1% osmium tetroxide (TAAB Laboratories Equipment Ltd.) in 0.1 M sodium cacodylate (Sigma-Aldrich) and 1% thiocarbohydrazide (Sigma-Aldrich) in ddH_2_O. Ears were dehydrated through a graded ethanol (Fisher Scientific) series (25% to 100%) at 4°C and stored in 100% acetone (VWR Chemicals) until critical point drying with liquid CO2 using an Emitech K850 (EM Technologies Ltd). Ears were mounted onto stubs using silver paint (Agar Scientific), sputter coated with palladium using a Quorum Q150R S sputter coater (Quorum Technologies) and visualised with a JSM-6010LV Scanning Electron Microscope (JEOL). Micrographs were pseudo-coloured in Adobe Photoshop.

## Results

### Identification of *CLRN2* as a novel deafness gene in a consanguineous Iranian family exhibiting autosomal recessive non-syndromic sensorineural hearing loss

A three generation Iranian family of Lurs ethnicity was ascertained as part of a large ethnically diverse Iranian population rare disease study (Fig. 1A). Three individuals that included the proband (IV-6), his sibling (IV-1), and a cousin (IV-8), born form consanguineous marriages, have reported moderate-to-profound bilateral non-syndromic sensorineural hearing loss (Fig. 1B). The age of onset for these three individuals was between 2 and 3 years of age. Pure-tone air-and bone-conduction audiometry thresholds (Fig. 1B) show evidence of intrafamilial variability. Individual IV-1 has a down sloping audiogram, with bilateral moderate-to-profound deafness. Individual IV-6 presented a moderate-to-severe hearing loss with slightly better hearing at higher frequencies. Both individuals showed normal (type A) tympanograms bilaterally. Speech recognition thresholds for individual IV-1 were 80 dB and 75 dB at 84% and 88% for right and left ears, respectively, and a most comfortable level of 95 dB. Speech recognition thresholds for individual IV-6 were 75 dB and 80 dB, each at 84%, for right and left ears, respectively. Patients have normal neuromotor, speech and language development, and did not show signs of impaired balance. No other abnormalities, including potential vision deficit, were present in the affected individuals, who were last evaluated at the age of 29 (IV-1), 44 (IV-6), and 25 (IV-8) years. For comparison, pure-tone audiometry was also recorded from a family member (V-1), with no reported history of hearing problem.

To identify the underlying genetic lesion, we applied homozygosity mapping in the extended family to identify a 15.2 Mb locus on chromosome 4p15.32p15.1 (GRCh37/hg19, chr4:17,298,445-32,495,165), defined by the SNPs rs7692897 and rs17081424 (Fig. 1C, Fig. S1A, Table S2). This locus contains 30 genes, none of which are presently associated with deafness in humans (Table S3). This approach also revealed four much smaller homozygous intervals on chromosomes 2p21 (137.3 kb), 3p22.2 (262.5 kb), 13q13.1 (90.7 kb), and 17q21.31 (292.6 kb) (Fig. S1A, Table S2) that do not contain known deafness-associated genes (Table S3). Pathogenic copy number variations were excluded. Next, we undertook exome sequencing of affected individual IV-6 (arrow, Fig. 1A). This generated 56,387,543 mappable reads, with 75.5% on-target reads. The mean depth was 57.3-fold, with 97.3% of regions with a 10-fold read depth. Analysis of the exome data of individual IV-6 excluded any candidate pathogenic variants in known deafness-associated genes (*42*) prompting an exome-wide analysis followed by filtering and re-analysis of variants in homozygous intervals (Table S4). Further, close inspection of the exome sequencing data revealed complete sequencing coverage of genes in the homozygous intervals (Table S5). Variant filtering detected a homozygous missense variant in *CLRN2* c.494C>A, (p.(Thr165Lys)) (NM_001079827.2) in the 15.2 Mb homozygous interval on chromosome 4 (Fig. S1A, S1B). This variant was shared with individual IV-1 and segregated in the extended family comprising a total of 14 individuals (Fig. 1A, D). Only individuals homozygous for the *CLRN2* c.494C>A variant exhibit hearing loss confirming the recessive nature of the allele (Fig. 1A).

### The *CLRN2* c.494C>A leads to a pathogenic missense substitution and aberrant splicing

The c.494C>A variant on chromosome 4p15.32 is unanimously predicted to be deleterious and disease causing by *in silico* tools (Table S6). The c.494C>A variant in *CLRN2* replaces a polar uncharged amino acid (threonine) with a positively charged amino acid (lysine) in clarin 2, (p.(Thr165Lys)) (*43*). This variant, as well as homozygous loss-of-function alleles are absent in population frequency databases. This suggests *CLRN2* is intolerant to biallelic loss-of-function. Our in-house collection of 89,041 additional exomes/genomes, including a multiethnic cohort of 842 exomes from probands with autosomal recessive hearing loss, identified four individuals from three families of Iranian, Turkish, and Emirati ethnicities, who carried the *CLRN2* c.494C>A variant (allele frequency 2.24×10^−5^). An Iranian hearing impaired individual was included among the carriers.

The c.494C>A variant involves the exchange of a novel polar threonine (Thr) residue to a basic lysine (Lys) amino acid that affects a highly conserved amino acid in the alpha-helix of the PMP-22/EMP/MP20/Claudin superfamily domain (Fig. 2A-C). Among clarin proteins, clarin 2 and clarin 1 show 34.9% identity with 81 identical and 91 similar amino acids (using UniProt (*44*), Fig. 2B). The outcome of *CLRN1* pathogenic or likely pathogenic missense variants, as well as nonsense variants (queried from the Deafness Variation Database v8.2 (*8*)) are marked in red (Fig. 2B) along with the clarin 2 p.(Thr165Lys) amino acid substitution (Fig. 2B, asterisk). Interestingly, nine out of the 19 clarin 1 amino acid mutated residues are identical in clarin 2. Three clarin 1 amino acid substitutions (p.(Leu163Pro), p.(Leu167Trp), and p.(Ile181Asn), NP_001182723.1) align in close proximity to the clarin 2 p.(Thr165Lys). Furthermore, clarin 1 p.Leu163Pro (45) and p.Ile181Asn (*46*), that are both reported in USH3A, are p.Leu150 and p.Ile168 in clarin 2. Most importantly, the threonine residue at position 165 (Thr165) CLRN2 is conserved across species and the corresponding amino acid in clarin 1 is a serine residue (Fig. 2A,B), a scenario often associated with conserved phosphorylation site residue, here by serine/threonine protein kinases (*43*).

**Fig. 2.**
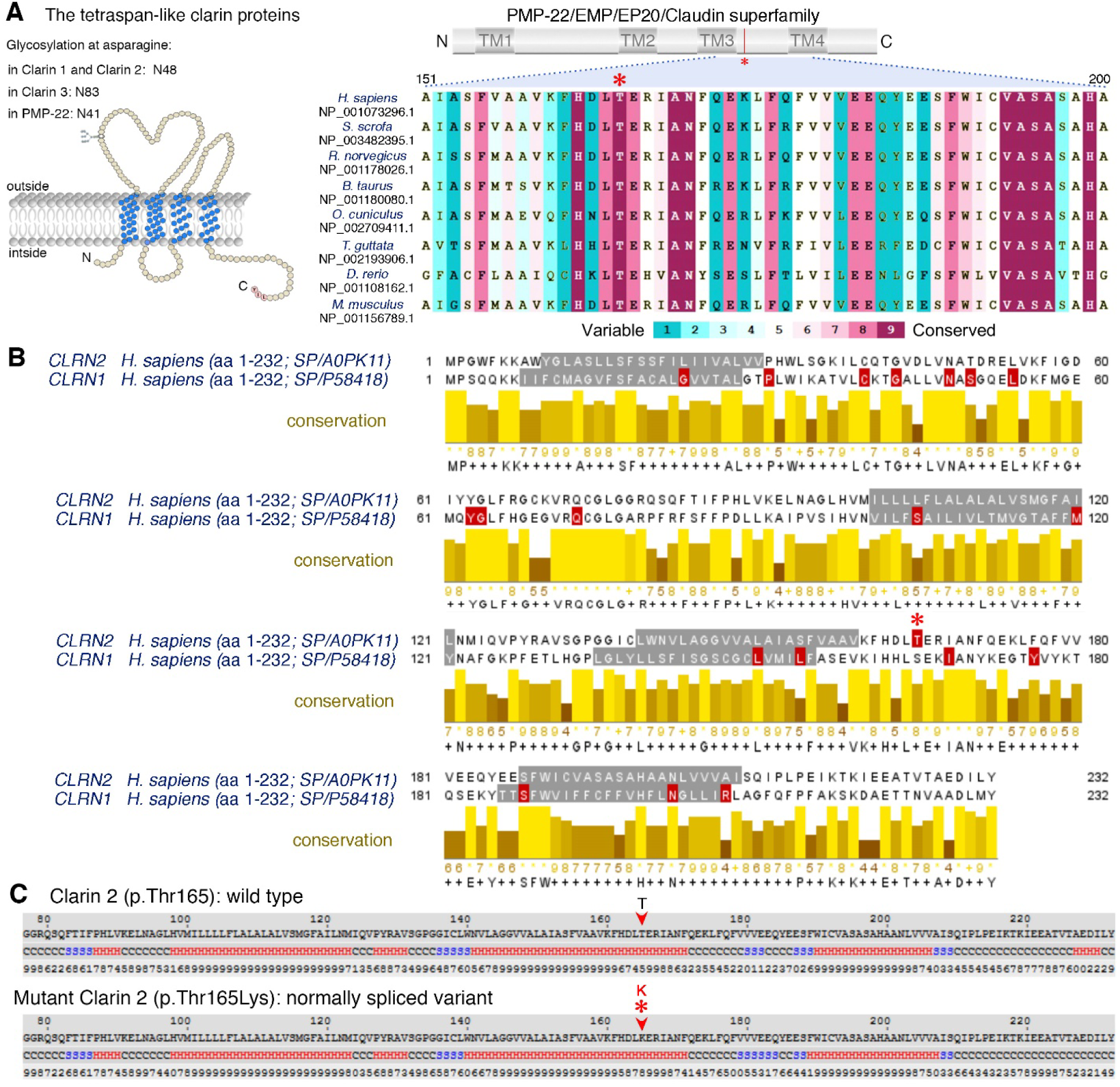
Conservation of the p.Thr165 residue, and clarin 1/clarin 2 alignment. **(A)** Overview of clarin 2 protein and modular structure of the PMP-22/EMP/EP20/Claudin superfamily, with amino acid residue coordinates and position of the p.(Thr165Lys) substitution shown (upper panel). An alignment of the amino acid sequences from the segment of clarin 2 (represented by dashed lines) from vertebrate species shows the Thr165 position (asterisk) is well conserved among vertebrates. **(B)** Alignment of clarin 2 (UniProtKB: A0PK11, upper alignment) and clarin 1 (UniProtKB: P58418, lower alignment) amino acid residues. Transmembrane domains are marked in grey, conservation is shown in yellow, and consensus sequences are shown below for the 232 amino acid proteins. Missense and nonsense variants in clarin 1 (Deafness Variation Database v8.2) and clarin 2 (present study, asterisk) are marked in red. **(C)** The predicted secondary structure of human clarin 2 (NP_001073296.1) wild-type (Thr165) and mutated (Thr165Lys) protein. H represents alpha-helix, S represents beta-strand and C represents coil.

In addition to causing an amino acid missense substitution, computational analysis also predicts that the c.494C>A variant will create an exonic splicing enhancer (ESE) motif, modifying the ESE hexameric sequence landscape of exon 3, which could interfere with the normal processing of *CLRN2* mRNA (Figs. S2A, S3B; ESEfinder and RESCUE-ESE, Human Splicing Finder) (*26, 27, 47*). To investigate the effect of the c.494C>A variant on *CLRN2* splicing, we used mini-gene assays using two different exon-trapping vectors and three different cell lines, Cos-7, ARPE-19, and HEK 293T. The mini-gene contained the 3’ end of intron 2, all of exon 3 (with and without the *CLRN2* variant), and ∼50 bp of the 3’ UTR (Fig. 3A) and was transfected into COS-7 and ARPE-19 cells. As a negative *CLRN2* control, we used the rs117875715 SNP, a common polymorphism, with a global minor allele frequency of ∼1.25% and >100 homozygous alleles reported in gnomAD (*23*) (http://gnomad.broadinstitute.org/variant/4-17528480-G-A) that is 20 nucleotides away from c.494C>A. Given its frequency, rs117875715 is predicted to be benign for hearing loss. Of note, this polymorphism is absent in the proband and family members reported here. Since exon 3 is the last exon of *CLRN2*, we designed our PCR primers to exclude the human poly-A signal and used the poly-A signal native to the pET01 vector. As expected for WT *CLRN2* (c.494C), we detected the splicing of the 5’ native pET01 exon only to exon 3 of *CLRN2* (Fig. 3A, B). The same normal splicing was obtained in all cell types transfected with *CLRN2* containing the control (rs117875715) variant (Fig. 3B). However, the c.494C>A variant yielded two bands; one ∼650 bp band matching the expected normally spliced exon, and a second abnormal band that was approximately ∼1,360 bp (Fig. 3B). Sequencing of these amplicons validated normal splicing including the c.494A variant and also revealed a retained intron 2 in the aberrantly spliced transcript (Fig. S3C). The retention of intron 2 results in a new reading frame that introduces a stop codon 26 amino acids after the native exon 2 splice site (p.(Gly146Lysfs*26)) (Fig. 3C). These results were replicated using the pSPL3b vector and HEK 293T cells (Fig. S3A-C), confirming the c.494C>A induced normal and aberrant splicing, independent of the cell type context. Following TA-cloning of cDNA amplicons from the homozygous individual (from Fig. S3B), 23 of 26 amplicons (88.5%) showed normal splicing, and 3 of 26 amplicons (11.5%) showed a retained intron.

**Fig. 3.**
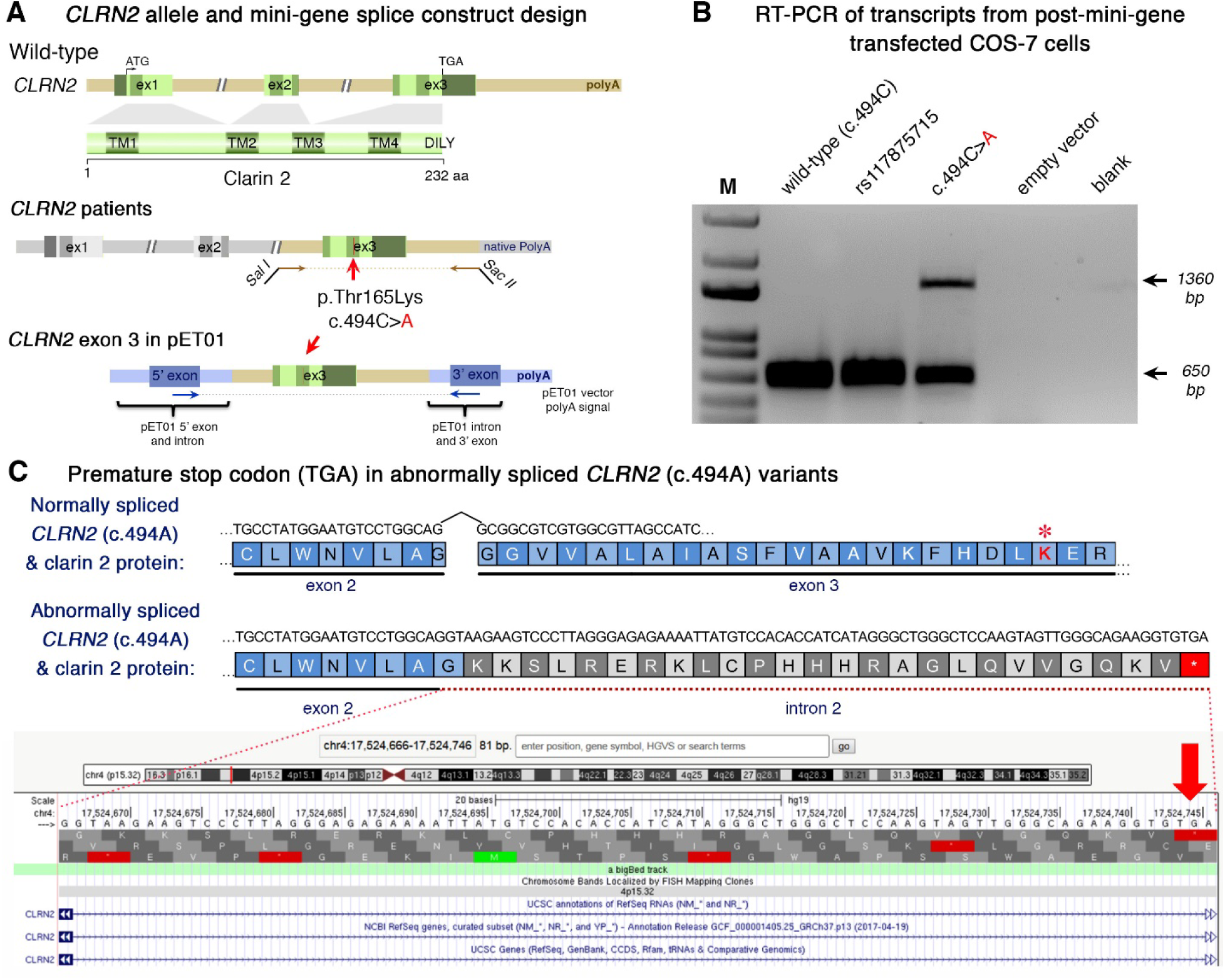
Analysis of the *CLRN2* c.494C>A variant on splicing. **(A)** Schematic illustration of the mini-gene splice construct design. Genomic representation of *CLRN2*, including the position of the missense variant c.494C>A (arrow) on exon 3 with 3’ UTR (green), and the 5’ UTR, as well as exons 1 and 2 (grey) (upper panel). Regions captured by mini-gene PCR primers are represented in green. Schematic illustration of the mini-gene splice construct including exon 3 and its flanking sequence (green) cloned into multiple cloning sites (*Sal*I and *Sac*II sites) of pET01 backbone vector (lower panel). Blue boxes represent native exons of the pET01 vector. **(B)** RT-PCR of transcripts from post-mini-gene transfected COS-7 cells. Amplicons derived from the transcripts of WT (*CLRN2*), a benign *CLRN2* polymorphism (rs117875715, chr4(GRCh37):g.17,528,480G>A), the *CLRN2* c.494C>A variant and a negative control, were visualized on a 1.5% agarose gel. The SNP, rs117875715, was used to test and validate the designed WT and mutant mini-gene assay. The ∼650 bp amplicon was associated with the WT and validation control rs117875715. The amplicon derived from the *CLRN2* c.494C>A transcripts showed two bands: a 650 bp band and a larger ∼1360 bp band, indicating retention of intron separating the donor site of the 5’ exon and the acceptor site of *CLRN2* exon 3. **(C)** Retention of intron in *CLRN2* c.494C>A mini-gene results in a stop codon (TGA) after *CLRN2* exon 2.

The mini-gene splicing assays and sequence analyses clearly show that the c.494C>A affects a highly conserved and key residue in clarin 2 sequence, while also creating aberrant mRNA splicing *in vivo* likely leading to a truncated protein. Altogether, this further confirms that variants in *CLRN2* can lead to sensorineural hearing loss.

### *Clrn2*, a hair cell expressed gene key to hearing also in zebrafish and mice

To further study the role of clarin 2 in the inner ear, we investigated its expression and analyzed potential impact of *Clrn2* loss-of-function in two other species, zebrafish and mice.

### clrn2 in zebrafish

Taking advantage of larva transparency, we used zebrafish as a model to investigate the *clarin 2* expression during early embryonic development. The RT-qPCR at different developmental stages revealed that *clrn2* mRNA was first detected at 18 hpf (Fig. 4A), a stage when the otic placode begins to form the otic vesicle in zebrafish (this stage is similar to mouse embryonic day 9 (E9), a stage of otic placode formation) (*48, 49*). *clrn2* mRNA expression increased (2-fold at 72 and 96 hpf compared to 18 hpf) and was maintained at later stages, up to 120 hpf (Fig. 4A). Comparative analyses of *clrn2* mRNA expression in different adult tissues of zebrafish revealed a significant enrichment in utricle, saccule and lagena of the inner ear (Fig. 4B). Our data are in agreement with RNA expression data from the Genotype-Tissue Expression (GTEx) project, wherein *CLRN2* mRNA in humans is enriched in the nervous system, testis, kidney, salivary gland, and lung. *CLRN1* has a similar expression profile in humans.

**Fig. 4.**
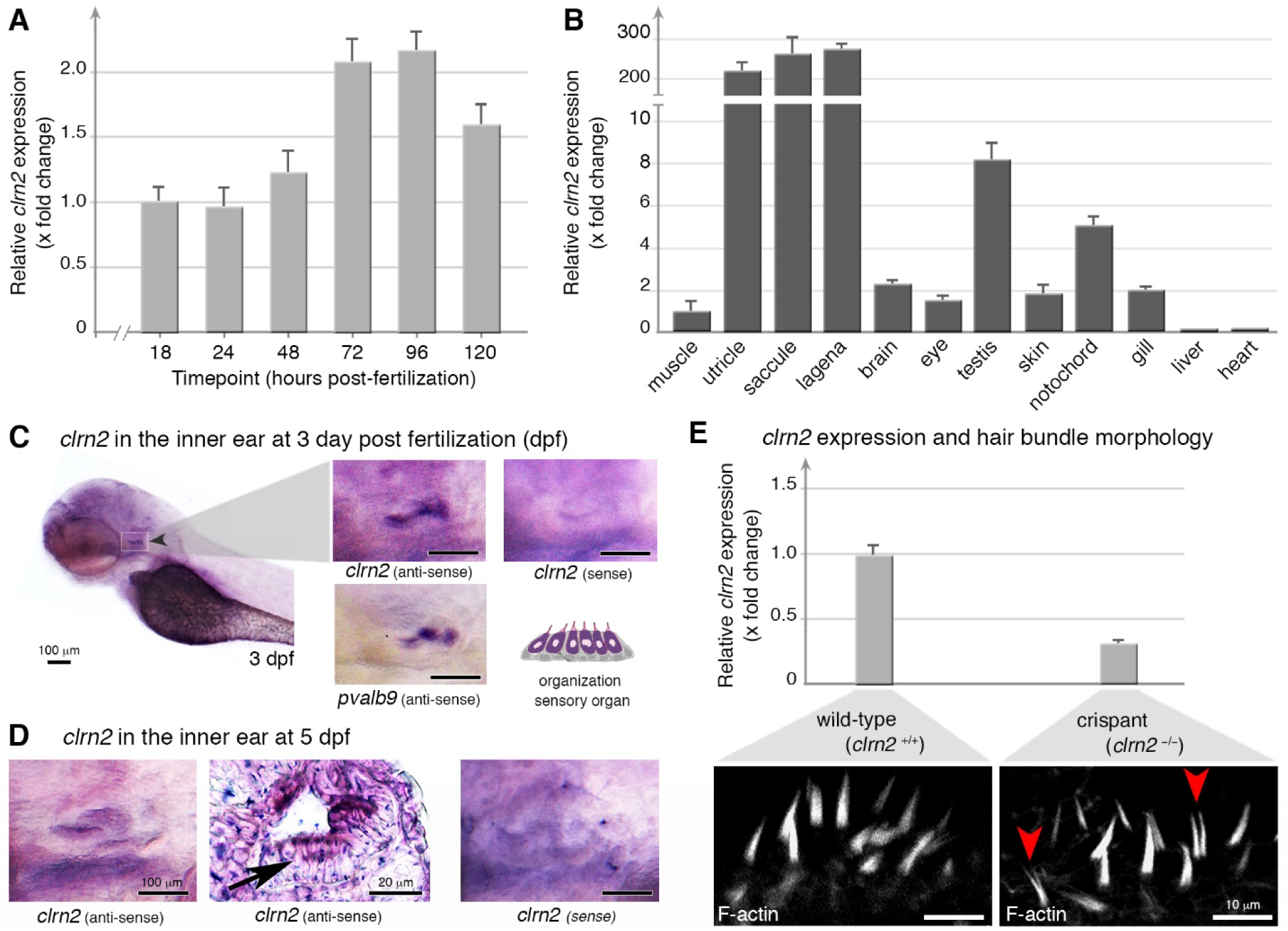
Clarin 2 is required for the inner ear function in zebrafish. **(A)** RT-qPCR of *clrn2* mRNA expression from 1 to 120 hpf of WT embryos/larvae. *clarin 2* mRNA expression can be detected starting from 18 hpf and then increased throughout development. The bar graphs showed the mean values ± SEM after normalization to the housekeeping gene *18S* level and then compared to 18 hpf. **(B)** RT-qPCR of c*lrn2* mRNA expression in different adult tissues. The bar graphs displayed mean values ± SEM after normalization to the housekeeping gene *18S* level and then compared to muscle. **(C, D)** Whole-mount in situ hybridization (WISH) using antisense *clrn2* probe reveals the inner ear expression of *clrn2* mRNA (relative dark purple color, black arrowhead) at 3 (C) and 5 (D) dpf embryos. Sense *clrn2* probe was used as negative control and relative light purple color is considered as background. *clrn2* mRNA was consistently expressed in hair cells within inner ear macula (C, D) with lined and arrayed structure. A known hair cell marker *pvalb9* was used as an indicator for hair cells in the inner ear of 3 dpf embryos (C). Cryosection was performed after *clrn2* WISH at 5 dpf to confirm the small patch of signal on the macula is hair cells rather than supporting cells (D, black arrow middle panel). Scale bar = 100 μm, except middle panel in D (20 μm). **(E)** RT-qPCR of *clrn2* mRNA expression level was decreased 70% in *clrn2* crispants compared to uninjected larvae, indicating *clrn2* was successfully knocked out (E, upper panels). Phalloidin staining on *clrn2* crispants show that the hair cells in the inner ear macula display splayed, thin and split structures (red arrowhead in lower panels). Scale bar = 10 μm.

To determine *clrn2* cellular expression, we used WISH in the inner ear of 3-and 5-dpf embryos (Fig. 4C, D). Unlike the *clrn2* sense probe, the anti-sense c*lrn2* revealed strong expression in the otic vesicle, similar to the expression of anti-sense *pvalb9*, used as a marker of hair cells (Fig. 4C). Histological examination of 5 dpf embryos further confirmed that *clrn2* expression is more specifically, restricted to hair cells, and is not expressed in the supporting cells of the inner ear (Fig. 4D).

To elucidate the function of *clrn2* in zebrafish, we used CRISPR/Cas9 to generate loss-of-function alleles. To maximize the knockout efficiency, we used five sgRNAs targeting the first and second exon of *clrn2* gene (Fig. S4). Injected embryos (crispants) were sequenced and, as expected, a mix of alleles in the form of deletions ranging from 4 bp to 73 bp, as well as insertions spanning +1 to +11 bp were observed. The majority of the variants were frameshift that would most likely create a premature stop codon in the protein (Fig. S4). The RT-qPCR analyses on injected embryos showed that c*lrn2* crispants have a significantly reduced amount of *clrn2* mRNA (Fig. 4E), suggesting nonsense mediated decay, leading to disrupted clarin 2 protein function.

Considering the expression in hair cells (Fig. 4D), we investigated the mechanosensory structures of the hair cell bundle, which are important for hearing and balance function in zebrafish. Interestingly, fluorescent phalloidin staining of the hair bundles of the inner ear in *clrn2* crispants (n=10) showed disrupted hair bundle structure compared to the WT controls; the hair bundles are splayed, thin and split in *clrn2* crispants (arrowheads in Fig. 4E). This defective phenotype, suggesting a critical role in hair bundle structures, is similar to the hair bundles in zebrafish *clrn1* knockouts (*50*), the *orbiter* mutants (defective in *protocadherin 15* (*pcdh15*), a gene associated with human Usher syndrome 1F) (*51*) and *ush1c* morphants and *ush1c* mutants (*52*).

### Clrn2 in mice

To further assess the requirement of clarin 2 for auditory function in mammals, and assess further its role in auditory hair bundles, we extended our analyses to mouse. Consistent with expression data in zebrafish (Fig. 4A, C, D), single cell RNA-seq data available to visualize on the gEAR portal (*umgear*.*org*) show that in the mouse cochlear epithelium at postnatal day 1 (P1) and P7, *Clrn2* transcripts are almost exclusively detectable only in inner and outer hair cell populations (*53*) (see also Fig. S5). We utilized a CRISPR/Cas9-engineered *Clrn2* loss-of-function mouse mutant, in which exon 2 has been deleted (*Clrn2*^*del629*^) (Fig. 5A), and measured ABRs in P21 (± 1 day) mice in response to click and tone-burst stimuli.

**Figure 5.**
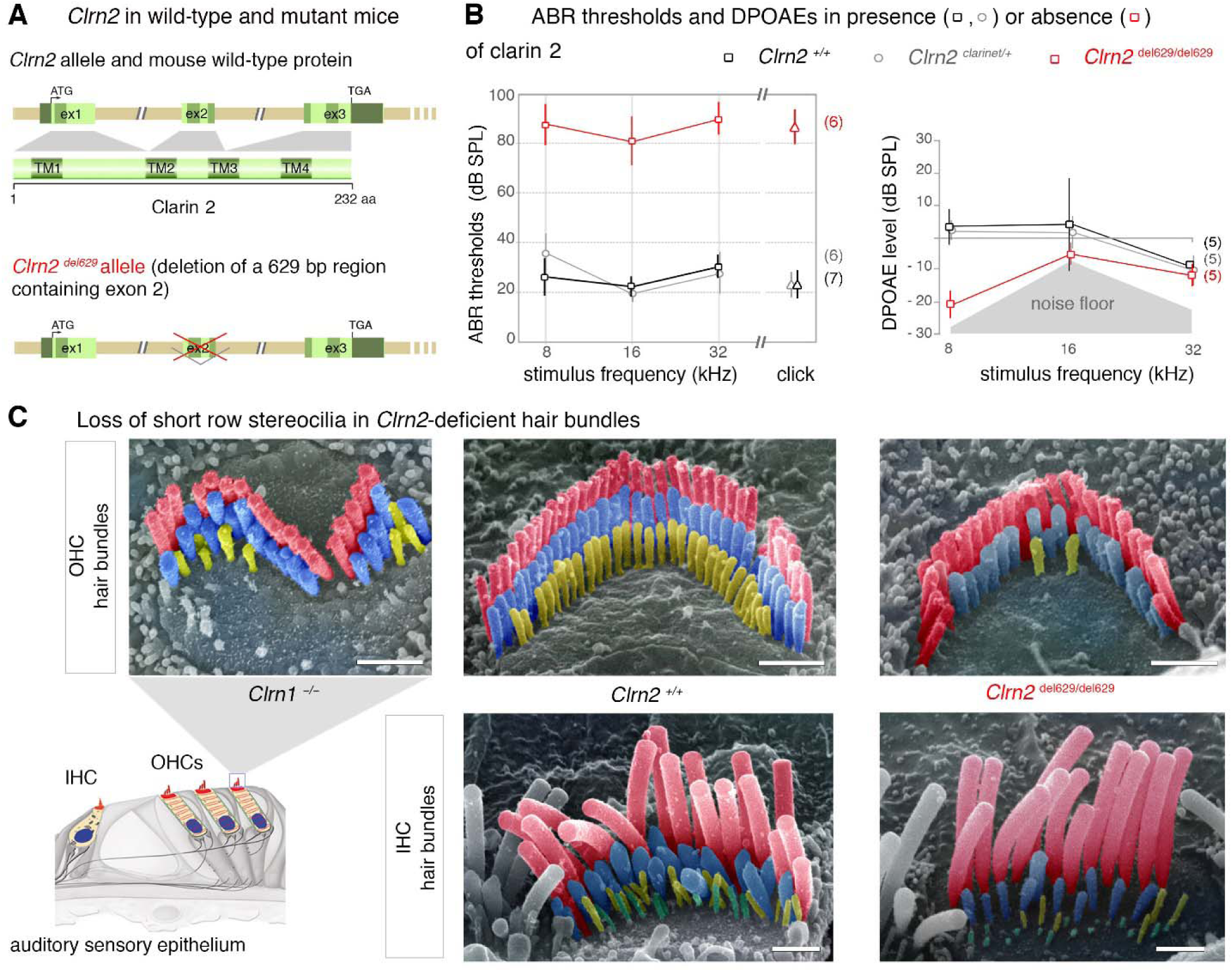
Clarin 2 is required for hearing function in mouse. **(A)** The genomic structure of mouse *Clrn2* (ENSMUST00000053250), and domains of the encoded tetraspan-like glycoprotein (232 amino acids). The positions of the transmembrane (TM) domains (dark green) and the structures of the WT *Clrn2 and Clrn2*^*del629*^ alleles are indicated. **(B)** ABR threshold measurements at P21 (± 1 day) show that *Clrn2* ^*del629/del629*^ mice (red) exhibit a severe-to-profound hearing loss affecting all frequencies tested, with thresholds at 80 dB SPL and beyond. Age-matched *Clrn2+/+* (black) and *Clrn2* ^*del629/+*^ (grey) controls display thresholds within the expected range (15-40 dB SPL). Data shown are mean ± SD. \*\*\**p*<0.001, one-way ANOVA. **(C)** Averaged DPOAE responses at P28 (± 1 day), showing significantly reduced responses in *Clrn2* ^*del629/del629*^ mice. Data shown are mean ± SD. \**p*<0.02, \*\**p*<0.01, one-way ANOVA. **(C)** Pseudo-colored scanning electron micrographs illustrate the three full rows, tallest (red), middle (blue) and short (yellow), of P28 (± 1 day) stereocilia in IHC and OHC hair bundles. Unlike the fragmented hair bundle in *Clrn1^−/−^* mice, lack of clarin-2 does not affect the shape of IHC or OHC hair bundles. However, all the short row stereocilia have completely or partially regressed in the absence of either clarin protein. Scale bar = 1 µm.

Analysis of ABR thresholds, which is the lowest sound stimulus required to elicit measurable activity in the auditory nerve, showed that homozygous (*Clrn2*^*del629/del629*^) mice display very elevated thresholds (>80 decibel sound pressure level (dB SPL)) at all frequencies tested: 8, 16 and 32 kHz (Fig. 5B). Whereas, *Clrn2*^*del629/+*^ mice exhibit thresholds comparable with those of WT (*Clrn2+/+*) littermates (<40 dB SPL), demonstrating the absence of a heterozygous auditory phenotype (Fig. 5B).

To further assess cochlear function, DPOAEs were measured in P28 (± 1 day) *Clrn2*^*del629/del629*^ mice. Compared to their *Clrn2+/+* and *Clrn2*^*del629/+*^ littermates, *Clrn2*^*del629/del629*^ mice have reduced DPOAEs (Fig. 5B) suggesting impaired outer hair cell (OHC) function.

To investigate stereocilia bundle morphology in *Clrn2*^*del629/del629*^ mice, we used scanning electron microscopy to examine the cochlear sensory epithelia. At P28 (± 1 day), the inner and outer hair cell stereocilia bundles of *Clrn2* mutant mice display the expected U-and V-shape, respectively, which contrasts with the grossly misshapen OHC bundles found in *Clrn1* mutant mice (Fig. 5C). However, while the patterning of the bundles appears normal in *Clrn2*^*del629/del629*^ mice the heights of their middle and short row stereocilia are visibly more variable compared with those of *Clrn2+/+* littermates, and many of the short row ‘mechanotransducing’ stereocilia are missing (Fig. 5C).

Together, our findings establish that clarin 2 is key to hearing function in zebrafish and mouse, and likely has a key role in the mechanotransducing stereocilia of the hair bundle.

## Discussion

We identify *CLRN2* as a novel deafness gene in human and zebrafish and describe a new deafness-causing allele in mice. Genetic study using homozygosity mapping and exome sequencing of an extended Iranian family with multiple consanguineous marriages identified a pathogenic variant, c.494C>A in exon 2 of *CLRN2* segregating with pre-lingual ARNSHL. The c.494C>A variant results in a missense and splicing defect in clarin 2. By producing mutant zebrafish and mice lacking clarin 2, we demonstrated the key role the protein plays to ensure normal structural and functional integrity of the hair bundle, the sound-and motion-receptive structure of inner ear hair cells.

The clarin gene family also includes the *CLRN1* gene. Pathogenic variants in *CLRN1* have been linked to variable clinical outcomes, ranging from non-syndromic RP (*11*) to USH3A characterized by variable and progressive post-lingual hearing loss, RP, and variable vestibular responses (*13*). Several cases of later onset HL and/or RP, as late as the sixth decade of life, have been reported for USH3A patients (*14*). Clinical examination of affected individuals in this family, at the age of 25 (IV-8), 29 (IV-1), and 44 (IV-6) years of age, excluded the presence of additional syndromic features showing that homozygosity for the c.494C>A variant causes non-syndromic hearing loss, ranging from moderate-severe (IV-6) to profound (IV-1) deafness (Fig. 1A, B).

The c.494C>A variant affects an amino acid that is highly conserved among PMP-22/EMP/EP20/Claudin superfamily proteins (Fig. 2A-C). In addition, the c.494 cytosine is highly conserved and the exchange to adenine is predicted to create an ESE site that likely impacts splicing efficiency in humans (Fig. S2A, B) but not zebrafish (Fig. S2C). We confirmed the effect on splicing using mini-gene assays. We showed that the c.494C>A variant acts in two ways: 1) as a missense variant (p.Thr165Lys) producing a mutant full length protein and 2) as a splice variant leading to intron retention (Fig. 3B, and Fig. S3B, C) expected to cause a premature stop codon 26 amino acids into intron 2 (p.Gly146Lysfs*26) (Fig. 3C).

Variants that disrupt splicing machinery signals can impact accurate recognition and removal of intronic sequences from pre-mRNA (*27*) and are recognized as significant contributors to human genetic diseases (*54*). ESE sequences are *cis*-acting elements primarily recognized by the SR family proteins that function by recruiting core splicing machinery components to splice sites or by acting antagonistically against nearby silencing elements (*27, 55, 56*). ESEs are often associated with introns that contain weak splicing signals, but they can also reside in exons and impact the splicing process.

Two potential mechanisms could synergistically contribute to the disruptive effect of the missense variant. *First*, the replacement of threonine with lysine, an amino acid with a positively charged ‘bulky’ side chain (lysine), may affect protein folding (*43*) and transport to the plasma membrane. Membrane proteins sort to the plasma membrane via the conventional secretory pathway associated with ER-to-Golgi complex (*57*). Misfolded membrane proteins are typically retained in the endoplasmic reticulum (ER) and degraded by the ER-associated degradation pathway (*58, 59*). It is possible that a small fraction of the misfolded clarin 2 p.(Thr165Lys) could reach the plasma membrane via the unconventional secretory pathway, similar to that reported for clarin 1 p.(Asn48Lys) (p.(N48K)) (*60*). The unconventional secretory pathway is induced by the ER–associated misfolded or unfolded protein response (*61, 62*). However, the mutant clarin 2 reaching the surface may be functionally inactive. *Second*, evolutionarily conserved threonine residues are also conserved protein phosphorylation sites. Phosphorylation adds a negative charge to the side chain of the amino acid and it serves as an important post-translational mechanism for regulation of protein function (*63*). Loss of threonine at position 165 would potentially prevent functional activation of clarin 2. However, additional experiments are essential to test these hypotheses and unravel the true pathogenic mechanism associated with the p.(Thr165Lys) missense variant.

Our *in situ* hybridization in zebrafish (Fig. 4C, D) and *in silico* analyses (Fig. S5) in mouse clearly support predominant expression of *Clrn2* in the sensory hair cells. To examine further the key role of clarin 2 in the inner ear, we generated zebrafish and mice lacking a functional protein. ABR measurements in *Clrn2*^*del629/del629*^ mice revealed an early-onset hearing loss with elevated hearing thresholds compared with their *Clrn2+/+* littermate controls (mean click threshold 87 dB SPL ± 7 s.d. and 24 dB SPL ± 6 s.d., respectively). These data are consistent with early-onset hearing loss observed in another loss-of-function *Clrn2* mutant (*Clrn2*^*clarinet*^), which harbors an early truncating nonsense variant (p.Trp4*) (*39*). Interestingly, unlike observations in humans with *CLRN1* pathogenic variants, neither *Clrn1* (*40, 64, 65*) nor *Clrn2* (*39*) knockout mice exhibit a retinal phenotype. In the inner ear, the reduced DPOAEs in both *Clrn2*^*del629/del629*^ and in *Clrn2*^*clarinet/clarinet*^ mice (*39*), indicates impairment of OHCs function. This, together with the severe-to-profound hearing loss already exhibited at P21 in *Clrn2* mutant mice points to gene defects likely affecting both inner hair cells (IHCs) and OHCs. This is further supported by scanning electron microscopy data showing loss of shortest row stereocilia in both the cochlear IHCs and OHCs (Fig. 5C). Phalloidin staining of *clrn2* crispants also confirms hair bundle abnormalities in zebrafish.

In regard to the observed progressivity of the hearing impairment in *clarinet* mice (*39*), the earliest reported clinical diagnosis of hearing loss of the *CLRN2* affected individuals in the family we present is between 2 and 3 years of age. Newborn hearing screening was not routinely performed when the affected individuals were born, so we cannot confirm hearing was normal at birth. In light of absent serial audiograms, we cannot report if the hearing loss experienced in these patients is progressive, as is observed in the mouse model (*39*).

In conclusion, we demonstrate the c.494C>A variant affects exon 3 splicing efficiency. We showed, for the first time, that *CLRN2* is a deafness-causing gene in humans. A variant causes hearing loss in humans, replicated by animal studies. Additional reports of families segregating *CLRN2* biallelic variants will be crucial to refine and dissect the clinical course and characteristics of hearing loss due to this gene. Together, our studies in zebrafish and mice establish that hearing loss is probably due to defective protein in the hair cells, where the presence of clarin 2 is essential for normal organization and maintenance of the mechanosensitive hair bundles.

## Acknowledgements

We would like to extend our gratitude to the family for their participation. We thank Dr. Caroline Lekszas, Dr. Daniel Liedtke, and Dr. Indrajit Nanda from the Institute of Human Genetics at the University of Würzburg for their technical expertise. This work was supported by Intramural Funding (*f*ortüne) at the University of Tübingen (2545-1-0 to B.V.), the Ministry of Science, Research and Art Baden-Württemberg (to B.V.), the Medical Research Council (MC_UP_1503/2 to M.R.B), ANR light4deaf (ANR-15-RHUS-0001), HearInNoise (ANR-17-CE16-0017), LHW-stiftung to A.E.), and a grant from NIH/COBRE GM103636 (Project 3); the Presbyterian Health Foundation (PHF) Grant to GKV. This study was funded in part by NIDCDs R01s DC002842 and DC012049 to RJS and T32 GM007748 to KTB. LAD is a Medical Research Council DPhil student (1774724).

