## Supplementary Figure 1-5; Supplementary Table 1-6 for "Biallelic mutation of *CLRN2* causes non-syndromic hearing loss in humans"

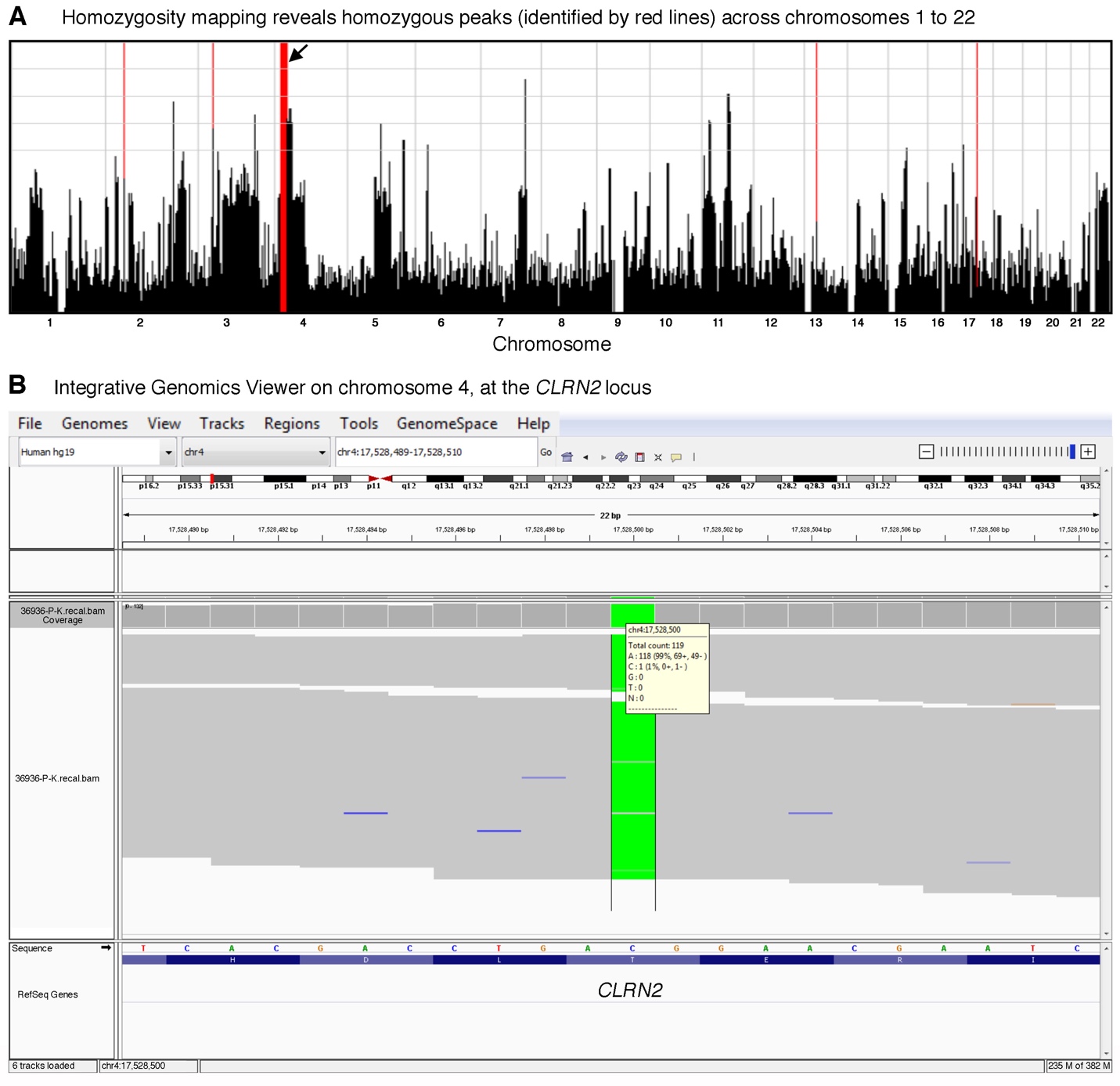

**Supplementary Figure 1. Homozygosity mapping profile and exome sequencing visualization of the *CLRN2* c.494C>A variant.**

**(A)** Homozygosity mapping showed homozygous peaks (identified by red lines) across chromosomes 1 to 22. An arrow shows the large homozygous stretch on chromosome 4. **(B)** Integrative Genomics Viewer visualization of the *CLRN2* c.494C>A putative pathogenic variant in individual IV-6.

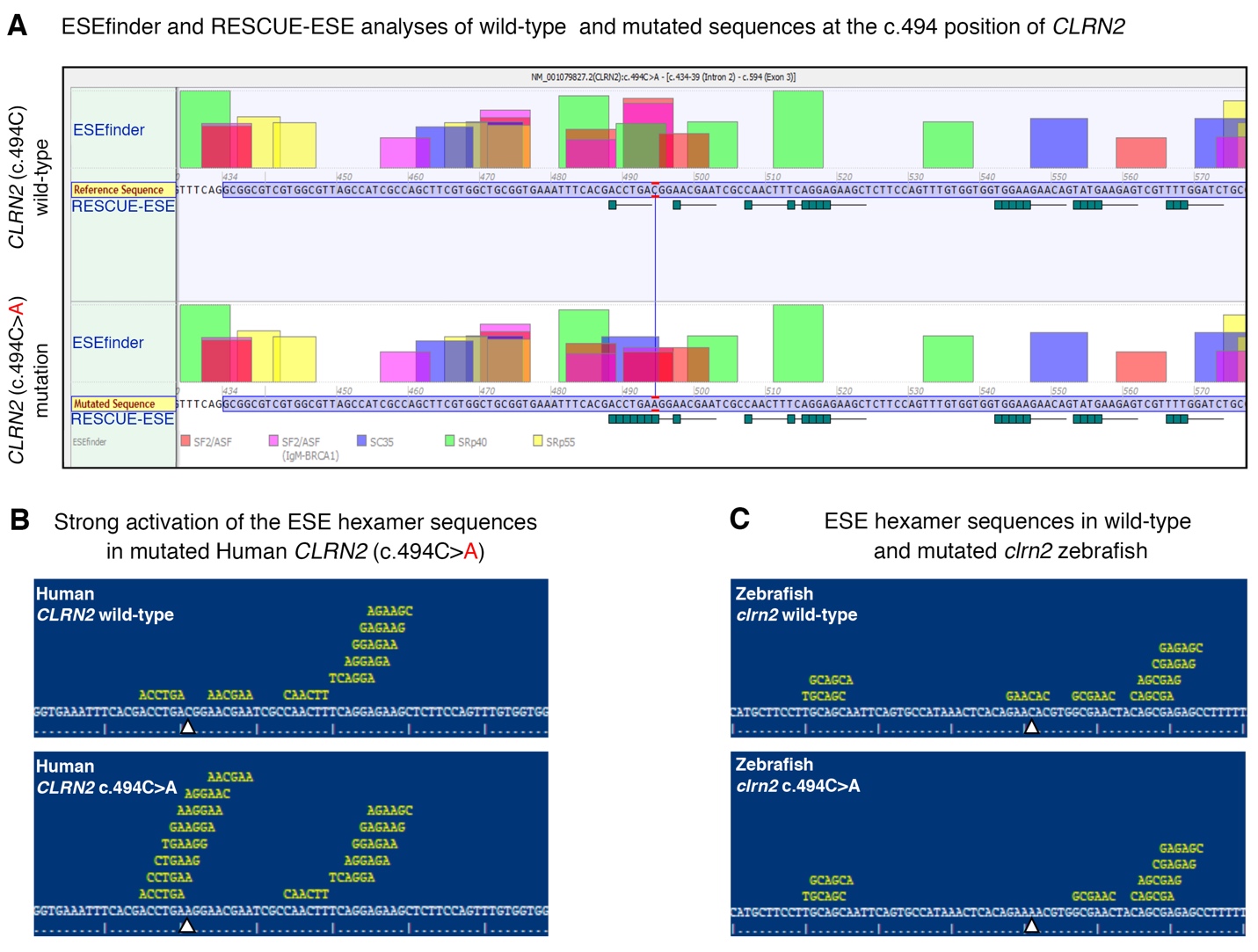

**Supplementary Figure 2. In silico prediction of the ESE splice site in *CLRN2*.**

**(A)** Analysis with ESEfinder and RESCUE-ESE reveal the splicing sequence landscape for the wild-type (upper panel) and mutated (lower panel) human sequence at c.494. The nucleotide at the c.494 position is outlined in red. ESE hits are displayed above each sequence. The green boxes represent RESCUE-ESE hexamers. The c.494C>T variant is predicted to induce an ESE hexamer that is shown by the string of green boxes in the bottom sub-panel. **(B)** Replication of the *CLRN2* splice prediction showing the wild-type (upper panel) ESE hexamer sequence (white arrow) compared to a strong activation of the ESE hexamer sequence (lower panel, arrow, increase in yellow hexameric sequence). **(C)** This hexameric sequence is not conserved in the zebrafish.

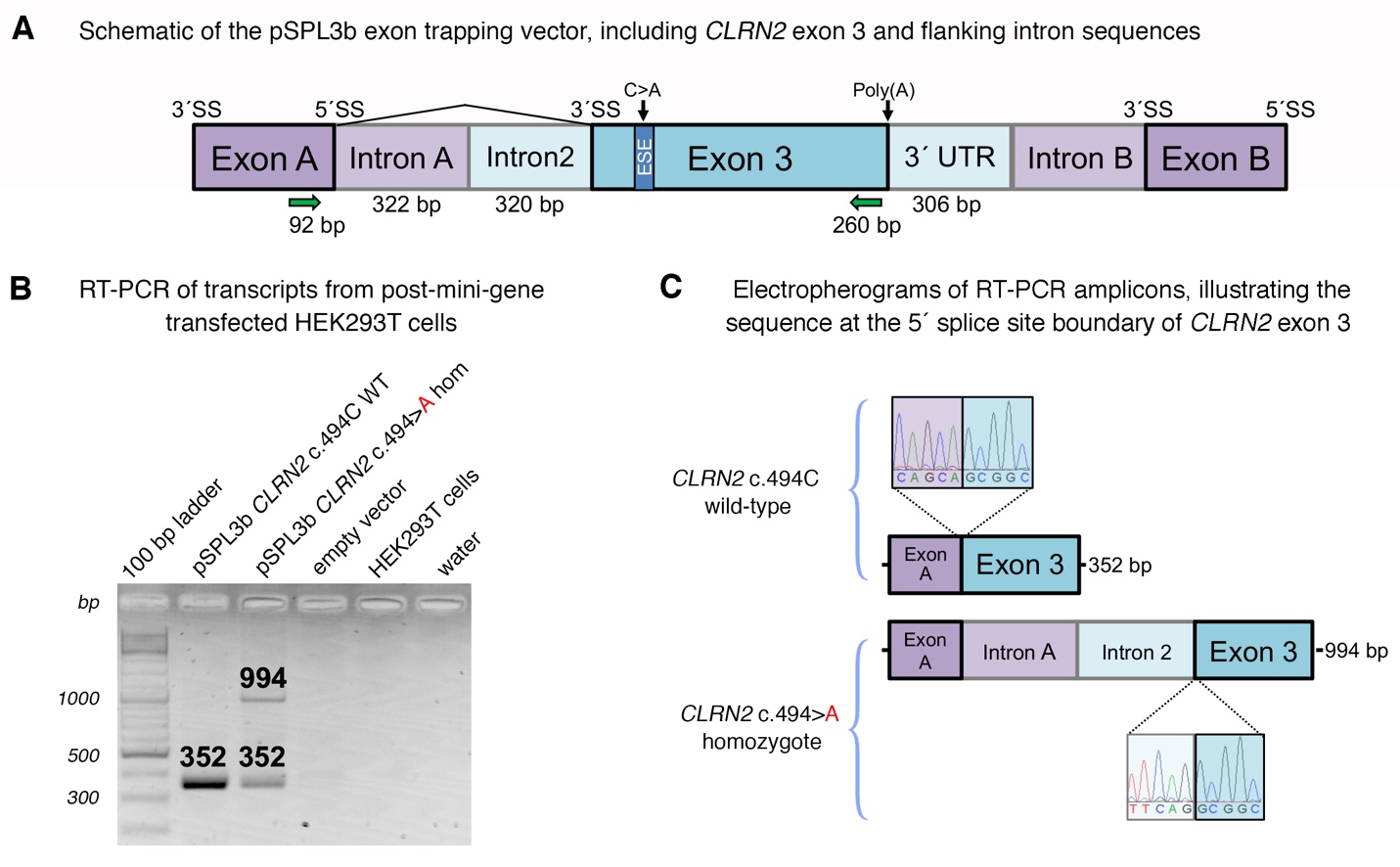

**Supplementary Figure 3. In vitro splicing assay of the *CLRN2* c.494C>A variant**

**(A)** Schematic of the pSPL3b exon trapping vector with cloned *CLRN2* exon 3 (medium blue) and flanking intron sequence (light blue) that was directly amplified from patient and wild-type family member genomic DNA. Exons A and B (medium purple) originate from the vector. 320 bp of intron 2 and 306 bp of the 3´ UTR were included (light blue). The C>A variant is predicted to create a exonic splice enhancer (ESE) in exon 3, represented by a blue bar. The primers that were used to amplify the Exon A splice donor (SD6) and *CLRN2* exon 3 splice acceptor regions are depicted by green arrows. **(B)** Electrophoretic visualization of cDNA RT-PCR products amplified from the constructs after transfection into HEK 293T cells. Amplicons were resolved on a 1.5% agarose gel. Wild-type splicing yields a 352 bp product that constitutes the Exon A and Exon 3 amplified regions. The mutant amplicon shows two bands, one of which shows a properly spliced 352 bp amplicon and the second band reveals a 994 bp amplicon that contains a retained exon trapping vector intron A and intron 2. **(C)** Electropheorgrams of the exon 3 5´ splice site boundaries for the RT-PCR products for wild-type (top) and mutant (bottom).

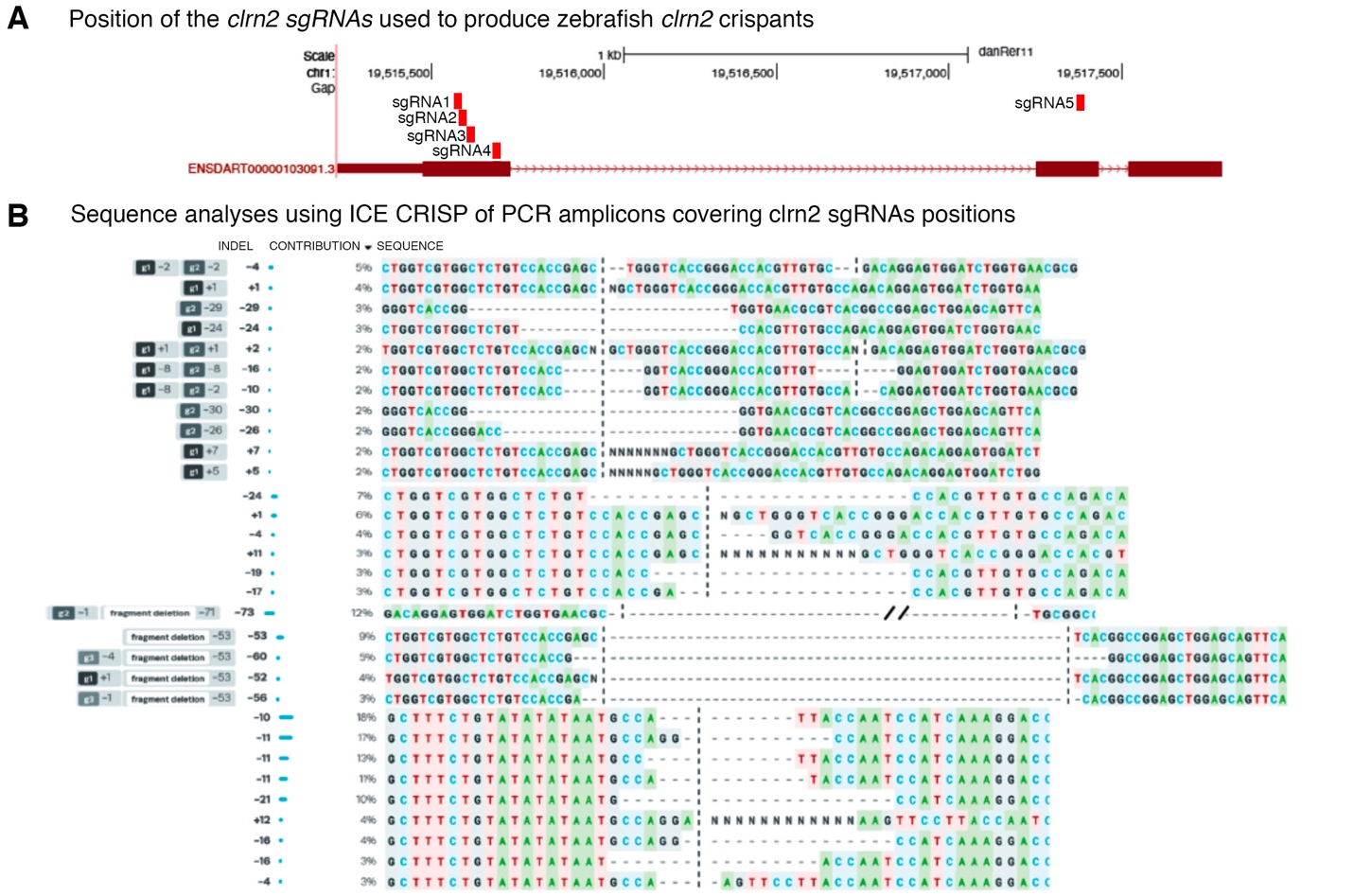

**Supplementary Figure 4. Generation of knockout alleles in the zebrafish *clrn2* gene using CRISPR/Cas9 mediated genome editing.**

**(A)** Five sgRNAs (red boxes) were designed in the *clarin-2* gene, four sgRNAs target exon 1 and one sgRNA targets exon 2. All five sgRNAs are mixed with Cas9 protein and injected together. **(B)** Mutations are identified using PCR and Sanger sequencing. A primer pair spanning sgRNA1-4 and sgRNA 5 was used to amplify 300 bp target region. Non-injected embryos were used as a control. PCR amplicons from a wild-type control and injected embryo were sequenced using Sanger sequencing. Sequence analysis was performed using ICE CRISPR analysis tools. Edited samples were aligned with the wild-type controls. All sgRNAs produced indels, and each embryo contains multiple small indels generating frameshift alleles, as well as large deletions between two sgRNAs.

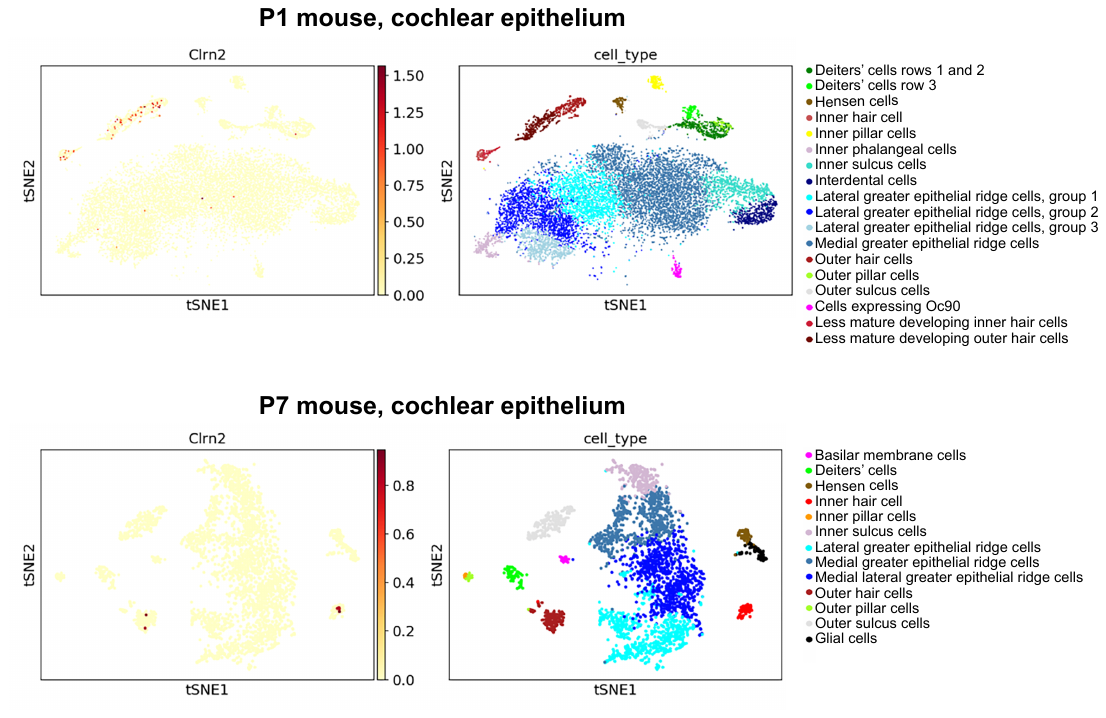

**Supplementary Figure 5. Visualization of single cell RNA-seq data from gEAR portal.**

The upper and lower panel shows tSNE plots of *Clrn2* expression in the cochlear epithelium at P1 and P7, respectively. Clustering is based on gene expression. Data^53^ were visualized on the gene Expression Analysis Resource (gEAR) (<https://umgear.org>).

**Supplementary Table 1. List of primers and sgRNA sequences used in the zebrafish study**

| Primer name | Sequence 5’→ 3’ |
| --- | --- |
| clarin-2-qPCR-forward | TCTTCACTGCTATTGGCTTTGC |
| clarin-2-qPCR-reverse | CATCGCACTAAACAGAGCTGC |
| 18S-qPCR-forward | AAACTGTTTCCCATCAACGAG |
| 18S-qPCR-reverse | GGGACTTAATCAACGCAAGC |
| clarin-2-BamHI-forward | agcGGATCCATGCCAACTCTCTGGAAGCAGA |
| clarin-2-XhoI-reverse | agcCTCGAGTCAGTAGAGGAAATCTTCTCCAGTGGC |
| clarin-2-sgRNA1 | taatacgactcactataGGTCCCGGTGACCCAGCGCTgttttagagctagaa |
| clarin-2-sgRNA2 | taatacgactcactataGGGACCACGTTGTGCCAGACgttttagagctagaa |
| clarin-2-sgRNA3 | taatacgactcactataGGATCTGGTGAACGCGTCACGGgttttagagctagaa |
| clarin-2-sgRNA4 | taatacgactcactataGGGGGCAAAACACGCAGATGgttttagagctagaa |
| clarin-2-sgRNA5 | taatacgactcactataGGATTGGTAAGGAACTTTCCgttttagagctagaa |
| Clarin-2_1_4-forward | TGAGCAGTGAAATGCCAACT |
| Clarin-2_1_4-reverse | CAAACATTCAAAACAATGTTGC |
| Clarin-2_1_5-forward | TTTTCCCAAAACTCATCAAAAA |
| Clarin-2_1_5-reverse | CATCGCACTAAACAGAGCTGA |

**Supplementary Table 2. Summary of common runs of homozygosity**

| Chromosome | Position start | Position end | rs start | rs end | Size (bp) |
| --- | --- | --- | --- | --- | --- |
| 2 | 47,612,878 | 47,750,227 | rs3924917 | rs6744097 | 137,349 |
| 3 | 37,034,946 | 37,297,443 | rs1800734 | rs11714766 | 262,497 |
| 4 | 17,298,445 | 32,495,165 | rs7692897 | rs17081424 | 15,196,720 |
| 13 | 32,888,473 | 32,979,138 | rs206115 | rs693963 | 90,665 |
| 17 | 41,152,503 | 41,445,145 | rs4028324 | rs9912203 | 292,642 |

**Supplementary Table 3. Gene content of homozygous intervals**

| Interval | Gene content |
| --- | --- |
| chr2:47,612,878-47,750,227 | *EPCAM*, *MSH2*, *KCNK12* |
| chr3:37,034,946-37,297,443 | *MLH1*, *LRRFIP2*, *GOLGA4* |
| chr4:17,298,445-32,495,165 | *QDPR*, *CLRN2*, *LAP3*, *MED28*, *FAM184B*, *DCAF16*, *NCAPG*, *LCORL*, *SLIT2*, *PACRGL*, *KCNIP4*, *ADGRA3*, *GBA3*, *PPARGC1A*, *DHX15*, *SOD3*, *CCDC149*, *LGI2*, *SEPSECS*, *PI4K2B*, *ZCCHC4*, *ANAPC4*, *SLC34A2*, *SEL1L3*, *SMIM20*, *RBPJ*, *CCKAR*, *TBC1D19*, *STIM2*, *PCDH7* |
| chr13:32,888,473-32,979,138 | *BRCA2*, *N4BP2L1* |
| chr17:41,152,503-41,445,145 | *RPL27*, *IFI35*, *VAT1*, *RND2*, *BRCA1*, *NBR1*, *TMEM106A* |

**Supplementary Table 4. Summary of variants in individual IV-6 after filtering settings were applied**

|  | Total variants remaining |
| --- | --- |
| Coverage ≥ 10-fold | 83,065 |
| Phred quality ≥ 30 | 27,660 |
| MAF ≤ 0.005 | 2,033 |
| Non-synonymous, indels, splice site | 579 |
| Removing artifact-prone genes^*^ | 332 |
| Homozygous | 46 |

^*^*HLA*s, *MAGE*s, *MUC*s, *NBPF*s, *OR*s, *PRAME*s

**Supplementary Table 5. Analysis of gene content in homozygous intervals for complete exome coverage, expression in the mouse inner ear, and mouse elevated ABR thresholds**

| Interval | Gene | Coordinates | Exome Coverage | Human OMIM phenotype | SHIELD Expression +/- | MGI Hearing Phenotype +/- | IMPC ABR Threshold |
| --- | --- | --- | --- | --- | --- | --- | --- |
| chr2:47,612,878-47,750,227 | | | | | | | |
| chr2 | *EPCAM* | chr2:47,596,287-47,614,167 | ✓ | Colorectal cancer, hereditary nonpolyposis, type 8, diarrhea 5, with tufting enteropathy, congenital | + | No info | No info |
| chr2 | *MSH2* | chr2:47,630,206-47,710,367 | ✓ | Colorectal cancer, hereditary nonpolyposis, type 1, mismatch repair cancer syndrome, Muir-Torre syndrome | + | No info | No info |
| chr2 | *KCNK12* | chr2:47,747,915-47,797,470 | ✓ | No human phenotype | + | No info | No info |
| chr3:37,034,946-37,297,443 | | | | | | | |
| chr3 | *MLH1* | chr3:37,034,841-37,092,337 | ✓ | Colorectal cancer, hereditary nonpolyposis, type 2, mismatch repair cancer syndrome, Muir-Torre syndrome | + | No info | Normal |
| chr3 | *LRRFIP2* | chr3:37,094,117-37,217,851 | ✓ | No human phenotype | + | No info | No info |
| chr3 | *GOLGA4* | chr3:37,284,682-37,408,370 | ✓ | No human phenotype | + | No info | No info |
| chr4:17,298,445-32,495,165 | | | | | | | |
| chr4 | *QDPR* | chr4:17,488,016-17,513,857 | ✓ | Hyperphenylalaninemia, BH4-deficient, C | + | No info | No info |
| chr4 | *CLRN2* | chr4:17,516,788-17,528,727 | ✓ | -- | + | No info | No info |
| chr4 | *LAP3* | chr4:17,578,927-17,609,590 | ✓ | No human phenotype | + | No info | No info |
| chr4 | *MED28* | chr4:17,616,273-17,626,160 | ✓ | No human phenotype | + | + | Abnormal |
| chr4 | *FAM184B* | chr4:17,633,709-17,783,135 | ✓ | -- | + | No info | Normal |
| chr4 | *DCAF16* | chr4:17,802,278-17,812,381 | ✓ | -- | No info | No info | No info |
| chr4 | *NCAPG* | chr4:17,812,436-17,846,487 | ✓ | No human phenotype | + | No info | No info |
| chr4 | *LCORL* | chr4:17,882,218-18,023,483 | ✓ | No human phenotype | + | No info | No info |
| chr4 | *SLIT2* | chr4:20,255,235-20,620,788 | ✓ | No human phenotype | + | No info | No info |
| chr4 | *PACRGL* | chr4:20,702,036-20,729,980 | ✓ | -- | + | No info | No info |
| chr4 | *KCNIP4* | chr4:20,730,239-21,699,318 | ✓ | No human phenotype | + | No info | Normal |
| chr4 | *ADGRA3* | chr4:22,388,997-22,517,677 | ✓ | No human phenotype | + | No info | No info |
| chr4 | *GBA3* | chr4:22,694,537-22,821,195 | ✓ | No human phenotype | No info | No info | No info |
| chr4 | *PPARGC1A* | chr4:23,793,644-23,891,700 | ✓ | No human phenotype | + | No info | No info |
| chr4 | *DHX15* | chr4:24,529,088-24,586,184 | ✓ | No human phenotype | + | No info | No info |
| chr4 | *SOD3* | chr4:24,797,085-24,802,467 | ✓ | Superoxide dismutase, elevated extracellular | + | No info | No info |
| chr4 | *CCDC149* | chr4:24,807,739-24,914,600 | ✓ | -- | + | No info | No info |
| chr4 | *LGI2* | chr4:25,000,471-25,032,414 | ✓ | No human phenotype | + | No info | No info |
| chr4 | *SEPSECS* | chr4:25,121,627-25,162,204 | ✓ | Pontocerebellar hypoplasia type 2D | + | No info | No info |
| chr4 | *PI4K2B* | chr4:25,235,653-25,280,831 | ✓ | No human phenotype | + | No info | No info |
| chr4 | *ZCCHC4* | chr4:25,314,396-25,372,005 | ✓ | No human phenotype | + | No info | No info |
| chr4 | *ANAPC4* | chr4:25,378,848-25,420,120 | ✓ | No human phenotype | + | No info | Normal |
| chr4 | *SLC34A2* | chr4:25,657,435-25,680,368 | ✓ | Pulmonary alveolar microlithiasis | + | No info | No info |
| chr4 | *SEL1L3* | chr4:25,749,049-25,864,610 | ✓ | -- | + | No info | No info |
| chr4 | *SMIM20* | chr4:25,915,814-25,931,501 | ✓ | No human phenotype | + | No info | No info |
| chr4 | *RBPJ* | chr4:26,322,429-26,436,752 | ✓ | Adams-Oliver syndrome 3 | + | No info | Normal |
| chr4 | *CCKAR* | chr4:26,483,018-26,492,042 | ✓ | -- | + | No info | No info |
| chr4 | *TBC1D19* | chr4:26,585,546-26,756,918 | ✓ | -- | + | No info | No info |
| chr4 | *STIM2* | chr4:26,862,313-27,027,003 | ✓ | No human phenotype | + | No info | Abnormal |
| chr4 | *PCDH7* | chr4:30,722,037-31,148,423 | ✓ | No human phenotype | + | No info | No info |
| chr13:32,888,473-32,979,138 | | | | | | | |
| chr13 | *BRCA2* | chr13:32,889,617-32,973,809 | ✓ | Fanconi anemia, complementation group D1, Wilms tumor, susceptibility to male breast cancer, familial breast-ovarian cancer 2, glioblastoma 3, medulloblastoma, pancreatic cancer 2, prostate cancer | + | No info | No info |
| chr13 | *N4BP2L1* | chr13:32,974,860-33,002,315 | ✓ | -- | + | No info | No info |
| chr17:41,152,503-41,445,145 | | | | | | | |
| chr17 | *RPL27* | chr17:41,150,446-41,154,971 | ✓ | ?Diamond-Blackfan anemia 16 | + | No info | No info |
| chr17 | *IFI35* | chr17:41,158,742-41,166,476 | ✓ | No human phenotype | + | No info | No info |
| chr17 | *VAT1* | chr17:41,166,622-41,174,459 | ✓ | No human phenotype | + | No info | No info |
| chr17 | *RND2* | chr17:41,177,258-41,184,058 | ✓ | No human phenotype | + | No info | No info |
| chr17 | *BRCA1* | chr17:41,196,312-41,277,500 | ✓ | Fanconi anemia, complementation group S, familial breast-ovarian cancer 1, susceptibility to pancreatic cancer 4 | + | No info | No info |
| chr17 | *NBR1* | chr17:41,323,246-41,363,707 | ✓ | No human phenotype | + | No info | - |
| chr17 | *TMEM106A* | chr17:41,363,894-41,371,589 | ✓ | No human phenotype | + | No info | No info |

-- not an OMIM gene

**Supplementary Table 6. Conservation and pathogenicity prediction scores for the *CLRN2* p.Thr165Lys variant**

| ***In-silico* prediction tool** | **p.Thr165Lys prediction score** |
| --- | --- |
| GERP++^1^ | Conserved (5.76) |
| PhyloP^2^ | Conserved (5.69) |
| CADD^3^ | 28.9 |
| LRT^4^ | Deleterious |
| MutationTaster^5^ | Disease causing (1) |
| PolyPhen-2^6^ | Probably damaging (0.995) |
| SIFT^7^ | Deleterious (0.0) |

^1^GERP++ scores > 0 are conserved, ^2^PhyloP ranges between -14.1 (not conserved) to 6.4 (conserved), ^3^CADD-20 represents a nucleotide variant in the top 1% of deleterious substitutions, CADD-30 represents a nucleotide variant in the top 0.1% of all deleterious substitutions, ^4^LRT scores range from unknown, neutral, and deleterious, ^5^MutationTaster scores closer to 1 represents a high probability of a damaging substitution, ^6^PolyPhen-2 scores between 0.85-1 represent a probably damaging outcome, ^7^SIFT scores between 0.0 and 0.05 represent a deleterious variant.
